# Sclerostin influences exercise-induced adaptations in body composition and white adipose tissue morphology in male mice

**DOI:** 10.1101/2022.06.29.498123

**Authors:** N Kurgan, J Stoikos, BJ Baranowski, J Yumol, R Dhaliwal, JB Sweezey-Munroe, VA Fajardo, W Gittings, REK MacPherson, P Klentrou

## Abstract

Sclerostin is an inhibitor of the osteogenic Wnt/β-catenin signalling pathway that has an endocrine role in regulating adipocyte differentiation and metabolism. Additionally, subcutaneous white adipose tissue (scWAT) sclerostin content decreases following exercise training (EXT). Therefore, we hypothesized that EXT-induced reductions in adipose tissue sclerostin may play a role in regulating adaptations in body composition and whole-body metabolism. To test this hypothesis, 10-week-old male C57BL/6J mice were either sedentary (SED) or performing 1h of treadmill running at ∼65-70% VO_2max_ 5 d/week (EXT) for 4 weeks and had subcutaneous (s.c) injections of either saline (C) or recombinant sclerostin (S) (0.1 mg/kg body mass) 5 d/week; thus, making 4 groups (SED-C, EXT-C, SED-S, and EXT-S; n=12/group). No differences in body mass were observed between experimental groups, while food intake was higher in EXT (p=0.03) and S (p=0.08) groups. There was a higher resting energy expenditure in all groups compared to SED-C. EXT-C had a higher lean mass and lower fat mass percentage compared to SED-C and SED-S. No differences in body composition were observed in either the SED-S or EXT-S groups. Lower scWAT (inguinal), vWAT (epididymal) mass, and scWAT adipocyte cell size and increased percentage of multilocular cells in scWAT were observed in the EXT-C group compared to SED-C, while lower vWAT was only observed in the EXT-S group. EXT mice had increased iWAT Lrp4 and mitochondrial content and sclerostin treatment only inhibited increased Lrp4 content with EXT. Together, these results provide evidence that reductions in resting sclerostin with exercise training may influence associated alterations in energy metabolism and body composition, particularly in scWAT.

## 1 Introduction

Sclerostin is a secreted glycoprotein found in the circulation, mainly produced by the osteocyte [1], that inhibits the canonical Wnt/β-catenin signalling pathway [2]. Within bone, Wnt/β-catenin signalling increases bone formation by increasing osteoblastogenesis. Thus, inhibiting sclerostin is a therapeutic target for treating osteoporosis [3]. In mice, genetic deletion, or inhibition of sclerostin with a neutralizing antibody, increases bone mass while also improving stimulated glucose uptake and lipid homeostasis [1, 4, 5]. Despite sclerostin not being expressed by adipose tissue (AT), reductions in AT mass, adipocyte cross-sectional area, and inhibited adipocyte maturation can also be observed in these models of sclerostin inhibition, ultimately conferring resistance to a high-fat obesogenic diet [1, 4, 5].

Given the recent identification of this novel endocrine function of sclerostin, the role it plays in regulating long-term adaptations to physiological stimuli has yet to be examined. Studies in humans support a physiological role of sclerostin in regulating metabolism in the context of exercise. Specifically, resting circulating sclerostin [6–10] and subcutaneous white AT (scWAT) content [11] have been shown to be reduced in humans following exercise training. Circulating sclerostin is also known to increase in response to acute exercise and the magnitude of this response is higher in adolescents with overweight or obesity (OW/OB) compared to normal-weight controls [12]. This suggests a bone-adipose tissue connection in response to exercise training through the regulation of peripheral sclerostin. Indeed, studies in rodents undergoing long term mechanical loading/exercise training show lower sclerostin expression and content within bone [13–16] and rodent models of obesity have higher expression of sclerostin within bone and higher resting serum sclerostin [17, 18]. Taken together, these results may explain the perturbed response of sclerostin to acute exercise between OW/OB and normal weight humans. Additionally, adipose tissue is reduced similarly in response to endurance exercise training [19, 20] and sclerostin inhibition [18] in rodents. These responses of peripheral sclerostin may influence tissue adaptations following exercise training through the regulation of Wnt signalling, particularly within scWAT [11, 21].

To test whether sclerostin is a regulator of exercise training-induced adaptations in body composition, the present study used continuous treatment of recombinant sclerostin to prevent potential exercise training-induced reductions in sclerostin. We aimed to examine if this would prevent subsequent adaptations in metabolism, bone mineral properties, body composition, or adipocyte cell size. We hypothesized that sclerostin may partially prevent exercise training induced reductions in AT mass and adipocyte cell size in response to exercise training through the regulation of Wnt/β-catenin signaling.

## 2 Materials and Methods

### 2.1 Animals

Experimental protocols complied with the Canadian Council on Animal Care and were approved by the Brock University Animal Care Committee (File #21-03-02). This study utilized one cohort of 48 male C57BL/6J mice (9 weeks of age) ordered from The Jackson Laboratory (Bar Harbor, ME, United States). Upon arrival, the mice acclimatized for 7d in the Brock University Comparative Biosciences Facility (St. Catharines, ON, Canada). Mice had *ab libitum* access to both water and food and were fed standard chow (2014 Teklad global 14% protein rodent maintenance diet, Harlan Tekland, Mississauga, ON, Canada). All mice were kept on a 12 h light / 12 h dark cycle.

### 2.2 Experimental Design

Baseline measurement of body weight and body composition with dual energy X-ray absorptiometry (DXA) of all mice (*n* = 48) was conducted on day 5 of acclimatization. In this study, mice were either sedentary (*n* = 24) or performing exercise training 5 d/wk (*n* = 24) and were further stratified to either treatment of 5 d/wk injections of recombinant sclerostin (0.1 mg/kg body mass) (cat# 1589-ST; R&D Systems, Minneapolis, Minnesota, USA) or saline control. Mice were randomized based on body fat percentage, measured with DXA (see below for details), into either one of four groups: Sedentary+Saline injections (SED+C) (*n* = 12), Exercise+Saline injections (EXT+C) (*n* = 12), Sedentary+Recombinant Sclerostin injections (SED+S), and Exercise+Recombinant Sclerostin injections (EXT+S). Body weight and food intake were monitored weekly. Exercise training was performed as previously described [22], which involved treadmill running for 1 h/d for 5 d/wk for 4 weeks. The speed and incline of the treadmill progressively increased from 20 m·min^-1^ at 10 ° for week 1 to 22 m·min^-1^ at 15 °, 23 m·min^-1^ at 20 °, and 25 m·min^-1^ at 20 ° for weeks 2 to 4, respectively. Mice were euthanized 48h after the final exercise bout and injections and inguinal white adipose tissue (iWAT), epididymal white adipose tissue (eWAT) depots, liver (only for necropsy tissue weight), serum, and tibias were collected. Necropsy tissue weights were also measured for WAT depots. One side of each WAT depot was taken for immunoblotting and the other for histology, and subsequently snap frozen in liquid nitrogen and stored at −80°C for future analysis.

### 2.3 Recombinant Sclerostin Injections

Mice either had subcutaneous (s.c) recombinant sclerostin (cat# 1589-ST; R&D Systems, Minneapolis, Minnesota, USA) or saline injections 5 d/week for 4 weeks, for a total of 20 s.c injections (insulin syringes (cat#: 01ST; ELIMEDICAL, Markham, Ontario, Canada) with 25G needles (cat#26403; EXELINT, California, USA) were used). The injection site was rotated on a day-to-day basis (e.g., Monday = scruff, Tuesday = right flank, Wednesday = left flank, Thursday = mid back, and Friday = low back) to reduce stress caused by repetitive injections to the same site. 0.01% Tween 80 and 1% Mannitol in phosphate-buffered saline (10 mM phosphate and 150 mM NaCl, pH 7.0) was used as the vehicle and the control. Calculating volume for injection was as follows: ((mouse mass (kg)*desired concentration (0.1 mg·kg body mass^-1^))/stock concentration (0.02 mg·ml^-1^)). For saline injected mice (control), volume was determined with the same calculation to control injection volume across groups. This concentration of recombinant sclerostin has been previously shown to increase circulating sclerostin immediately post-injection by 75% and by 20% 24h post-injection, which is a similar range of exercise training-induced decreases in sclerostin in humans [6–10].

### 2.4 Exercise Test to Exhaustion

All 48 mice ran on a 20° incline at an initial speed of 12 m·min^-1^ with speed increasing by 1 m·min^-1^ after 2 min, 5 min, and every additional 5 min of exercise up to 40 m·min^-1^ where they remained until exhaustion. However, 2 mice lacked motivation to exercise and were removed from analysis (one from SED-S and one from EXT-S). Exhaustion was determined by an inability to remain halfway up the treadmill before falling off the moving belt four times consecutively despite light encouragement (based on [23]). Time to exhaustion and distance travelled were monitored and used to assess work (Work (J) = body mass (kg)*gravity (9.81 m/sec^2^)*vertical speed (m/sec x angle)*time (sec)) and power (Power (W) = work (J)/time (sec)) [24].

### 2.5 Metabolic Caging

During the end of the final week of the intervention, a subset of 32 mice (n = 8/group) were single housed in metabolic cages (Sable Systems International, North Las Vegas, USA, Promethion high-definition behavioural phenotyping system) [25] secured in thermal cabinets set to 25°C with a 12h light/ 12h dark cycle schedule for 48 hours. Oxygen consumption (VO_2_), carbon dioxide expiration (VCO_2_), energy expenditure (kcal), respiratory exchange ratio, and total physical movement (i.e., activity in metres; measured by infrared beam breaks) were recorded continuously and averaged every 30 min using Sable Systems data acquisition software (IM-3 v20.0.6). Data were analysed using Sable Systems International MacroInterpreter software using One-Click Macro (V2.35.2). Mass-dependent variables (VO_2_, VCO_2_ and energy expenditure) were normalized to body weight. To examine differences in substrate utilization, VO_2_ and VCO_2_ were used to calculate carbohydrate and fat oxidation: carbohydrate oxidation= 4.585 VCO2 production (L·min^−1^) − 3.226 VO2 production (L·min^−1^); fat oxidation = 1.695 VO2 production (L·min^−1^) − 1.701 VCO2 production (L·min^−1^). Values were divided by 60 and multiplied by 16.19 (carbohydrate) and 40.80 (fat) to convert to kilojoules per hour (kJ/h)) [26].

### 2.6 Whole Body Composition and Bone Properties

Mice were anesthetized (2-3% isoflurane) and underwent whole body composition (lean mass (g), lean mass percentage, fat mass (g), and fat mass percentage), bone mineral density (BMD) (g·cm^-2^), bone mineral content (BMC) (g), and bone area (cm^2^) analysis using the iNSiGHT Dual Energy X-Ray Absorptiometry (DXA) system (Osteosys, Korea) during the week prior to the beginning of the study and following the fourth and final week. The inhouse coefficient of variation (CV) for the DXA is less than 2.5% for all measures, which was determined by the same operator imaging, readjusting, and analyzing the same mouse 6 times.

### 2.7 Trabecular and Cortical Bone Structure of the Tibia

Using micro-computed tomography (SkyScan 1176 V.1.1 (build 12), Bruker microCT, Belgium), *ex vivo* analysis of trabecular and cortical bone structure was conducted at the proximal tibia and tibia midpoint, respectively. Methods were conducted as previously described [27, 28]. Briefly, the isolated tibia was wrapped in parafilm wax and secured to the scanning bed within a Styrofoam holder. A 0.25 mm aluminum filter was applied to the high resolution (9 µm) scans. Under 850 ms of X-ray radiation exposure (45 kV voltage and 545 µA amperage), images were acquired at 0.2° increments over a 180° frame. Upon completion, image reconstruction (NRecon V.1.7.3.1 software, Bruker microCT, Belgium) was performed based on the following parameters: smoothing (3), ring artifact reduction (6), beam hardening (40), and a dynamic image range (0-0.148577). DataViewer V.1.5.6.2 (64-bit) software (Bruker microCT, Belgium) was then used for reorientation. Regions of interest were manually delineated for quantification (CT Analyzer V.1.17.7.2+ (64-bit), Bruker microCT, Belgium) of trabecular and cortical bone structure at the proximal tibia (offset: 0.450 mm from growth plate reference point; height: 0.675 mm) and tibia midpoint (height: 0.900 mm), respectively. Trabecular bone structure outcomes included bone volume fraction (BV/TV, %), trabecular thickness (Tb.Th, mm), trabecular separation (Tb.Sp, mm), trabecular number (Tb.N, 1/mm), degree of anisotropy (DA, no units), and connectivity density (Conn.Dn, mm^3); which were calculated using an adaptive thresholding of 49 (low) and 255 (high). Cortical bone structure outcomes included cortical area fraction (Ct.Ar/Tt.Ar, %), cortical thickness (Ct.Th, mm), periosteal perimeter (Ps.Pm, mm), endocortical perimeter (Ec.Pm, mm), medullary area (Ma.Ar, mm175:19542), and eccentricity (Ecc, no units); which were calculated using an adaptive thresholding of 74 (low) and 255 (high). One operator performed all imaging and analysis and was blinded to the study groups.

### 2.8 Insulin Tolerance Testing

Intraperitoneal insulin tolerance tests (ITT) were performed on non-anesthetized mice following 48h of no exercise. Tail vein was sampled for blood glucose using an automated glucometer (FreeStyle Libre) at baseline and at 15, 30, 45, and 60 min following an intraperitoneal injection (i.p.) of insulin (0.75 U·kg body mass^-1^) and area under the curve (AUC) of the glucose response over time was assessed.

### 2.9 Blood Sampling and ELISA

At end point, blood samples were taken by cardiac puncture with 1 mL insulin syringes (cat#: 01ST; ELIMEDICAL, Markham, Ontario, Canada) and 25G needles (cat#26403; EXELINT, California, USA). Blood samples were left on ice to clot for 1h before being centrifuged for 15 min at 1500 x g. Samples were aliquoted and stored at −80°C until analysis. Serum sclerostin was analysed in duplicate using a Quantikine enzyme-linked immunoassay kit (cat#MSST00; R&D Systems, Minneapolis, Minnesota, USA) according to manufacturer’s instructions.

### 2.10 Adipose Tissue Processing and Embedding

Following dissection, WAT depots were fixed in 10% neutral buffered formalin (cat#16004-126; VWR, Radnor, Pennsylvania, USA) for 48 h. Tissue samples were then stored in 70% ethanol for 24 h prior to histology. To initiate the embedding process, samples were placed in cassettes (cat#18000-136; VWR, Radnor, Pennsylvania, USA) and treated with 90% ethanol for 30 min on a stir plate and then treated with 3 passages of 100% ethanol for 45 min each. Next, tissues were treated with xylene (cat#UN1307; Thermo Fisher Scientific, Waltham, Massachusetts, USA) for 45 min on a stir plate, this step was repeated two more times, with samples being transferred to fresh xylene each time. Tissues were then placed in paraffin (cat#8330; Thermo Fisher Scientific, Waltham, Massachusetts, USA) for 60 min in a cell incubator at 57°C. Samples were then transferred to fresh paraffin for another 60 minutes in the incubator at 57°C. Subsequently, the samples were embedded in paraffin using the Paraffin Embedding Station (cat#EG1150H/EG1150C; Leica Biosystems, Wetzlar, Germany) and sectioned at 10 µm using the Rotary Microtome HM 325 (Thermo Fisher Scientific, Waltham, Massachusetts, USA). Sections were placed on a water bath set to 37°C and three sections were placed on a slide. Two slides were made for each representative sample. Slides were left to dry at room temperature overnight prior to proceeding to the staining.

### 2.11 Adipose Tissue Hematoxylin and Eosin Staining

Slides were deparaffinized in two 10 min treatments of xylene. The samples were then rehydrated in 95% ethanol for 2 min and 70% ethanol for 2 min. Slides were rinsed in distilled water and then placed in hematoxylin stain (cat#SH26-500D; Thermo Fisher Scientific, Waltham, Massachusetts, USA) for 3 min. Slides were then placed in tap water for 2 min and excess water was tapped off and slides were treated with a 0.2% acid-alcohol solution (100 ul of concentrated hydrochloric acid in 50 ml ethanol) for about 30 s. The slides were then placed in the Eosin Y (cat#SE23-500D; Thermo Fisher Scientific, Waltham, Massachusetts, USA) for 15-20 s and the excess stain was removed with distilled water. The slides were then dehydrated in 80% ethanol for 2 min, 90% ethanol for 2 min, and then 100% ethanol for 2 min. Following the dehydration treatment, slides were placed in xylene for 2 min and then cover slipped with Permount (cat#SP15-500; Thermo Fisher Scientific, Waltham, Massachusetts, USA). For image analysis, three representative images of the samples were taken with a Cytation 5 cell imaging multi-mode reader with Gen5 software (Version 3.10; Biotek, Vermont, USA) and analyzed using ImageJ. The mean surface area and percent multilocularity were measured. One researcher performed all imaging and analysis and were blinded to groups.

### 2.12 Adipose Tissue Homogenization

iWAT were homogenized (FastPrep®, MP Biomedicals, Santa Ana, CA) in 3x volume per mg weight of tissue of NP40 Cell Lysis Buffer (cat# FNN0021; Life Technologies) supplemented with 3x the recommended volume of phenylmethylsulfonyl fluoride (PMSF) and protease inhibitor (PI) cocktail (cat#7626-5G and cat#P274-1BIL, respectively; Sigma-Aldrich, St. Louis, Missouri, USA). Following homogenization, samples were placed on ice for 30 min then centrifuged at 4°C for 10 min at 5,000 x g. The infranatant was then collected and protein concentration was determined using a Bicinchoninic acid assay (cat#B9643; Sigma-Aldrich, St. Louis, Missouri, USA; cupper (II) sulfate pentahydrate, cat#BDH9312, VWR, Radnor, Pennsylvania, USA). The samples were prepared to contain equal concentrations (1 μg·μl^-1^) of protein in 1x Laemmli buffer (cat#1610747, Bio-Rad, Hercules, California, USA) and stored at −80°C for future analysis.

### 2.13 Immunoblotting

20 µg of protein were loaded and resolved on 10% TGX fast cast gels (cat#1610173, Bio-Rad, Hercules, California, USA) for 30 min at 250 V. Proteins were then either semi-dry transferred onto a polyvinylidene difluoride membrane at 1.3 A and 25 V for 7 min (Trans-Blot® Turbo™ Transfer System, Bio-Rad, Hercules, California, USA) or wet transferred onto nitrocellulose membrane at 100 V for 90 min (i.e., Lrp4). Membranes were blocked in Tris buffered saline/0.1% Tween 20 (TBST) with 5% non-fat powdered milk for 1h at room temperature. Primary antibody (1:500-2000 ratio dilution in 5% milk) was then applied and left to incubate on a shaker at 4°C for ∼16 h. Membranes were then washed with TBST 3 x 5 min and then incubated with the corresponding secondary antibody conjugated with horseradish peroxidase (anti rabbit (cat#HAF008), goat (cat#CAF109), and mouse IgG (cat#HAF007) – 1:2000 dilution in 5% milk, purchased from R&D Systems, Minneapolis, Minnesota, USA) for 1h at room temperature. Signals were detected using Clarity™ Western chemiluminescent substrate (cat#170-5061, Bio-Rad, Hercules, California, USA) and were imaged using a Bio-Rad ChemiDoc™ Imaging System (Hercules, California, USA). Each membrane was stained with Ponceau S to confirm equal protein loading and used to normalize. Densitometry analysis was done using ImageLab Software (Bio-Rad, Hercules, California, USA). Antibodies against total GSK3β (cat#12456), serine 9 phospho-GSK3β (cat#9336), active β-catenin (Ser33/37/Thr41) (cat#8814), total HSL (cat#4107), phosphor-HSL (563: cat#4137 and 660: cat#45804), and total ATGL (cat#2439) antibodies were purchased from Cell Signalling (Danvers, USA). Sclerostin (cat#MAB1406) and Lrp4 (cat#MAB5948) antibodies was purchased from R&D Systems, Inc. (Minneapolis, Minnesota, USA). Total oxphos antibody cocktail (cat#ab110413) and UCP1 (cat#ab10983) and serine 406 phospho-ATGL (cat# ab135093) antibodies were purchased from Abcam (Cambridge, United Kingdom). PGC-1α (cat#AB3242) was purchased from Millipore (Burlington, Massachusetts, USA).

### 2.14 Statistical Analysis

Comparisons of all factors were assessed with a 2-way ANOVA (Exercise and Injection) and significant interactions were followed up with pairwise comparisons with *Tukey* correction using GraphPad Prism 9 (San Diego, California, USA). Data are presented as means ± standard deviation (SD) with significance assumed at an α < 0.05.

## 3 Results

### 3.1 Sclerostin influences food intake and energy expenditure but does not further alter their responses to exercise training

The progressive exercise test to exhaustion following the intervention showed EXT mice were able to run longer (time) and obtain a higher top speed (resistance) than SED mice leading to greater work output (main effect for exercise p < 0.0001, grand mean difference = +3111 J) with no influence of sclerostin on this adaptation (exercise*injection interaction p = 0.3) and no effect in general (main effect for injection p = 0.7) (Figure 1a). Weekly food intake was higher in EXT mice compared to SED mice (main effect for exercise p = 0.03, grand mean difference = +1.85 g/wk). There was no influence of sclerostin on exercise training induced increases in food intake (exercise*injection interaction p = 0.6), but food intake trended to be higher in sclerostin injected mice compared to saline (main effect for injection p = 0.08, +1.46 g/wk) (Figure 1b). Body weight was also higher in EXT mice compared to SED mice (main effect for exercise p = 0.05, grand mean difference = +1.15 g). Despite a trending increase in food intake, there was no influence of sclerostin on body weight (main effect for injection p = 0.3) or on the EXT induced increase in body weight (exercise*injection interaction p = 0.4) (Figure 1c). Cage activity was not different between injection type (p = 0.7) or in exercise training mice (p = 0.09) during the dark cycle, but there was a simple effect for exercise training (p = 0.01, +28.0m) and an exercise*injection interaction (p = 0.01) during the light cycle that was a result of EXT-S mice being more active compared to SED-S mice during the light cycle (p = 0.01, +50.0m) while EXT-C mice had slightly higher cage activity (non-significant) compared to SED-C mice (p > 0.09, +6.0m). It is important to note that exercise training was done during the beginning of the light cycle (08:00-10:00), which could be why we didn’t see higher activity in the dark cycle compared to the light cycle in EXT mice. Despite no difference in cage activity between injection type that could influence energy expenditure, there was an exercise*injection interaction during the light (p = 0.0006) and dark cycles (p = 0.0005) for energy expenditure (Figures 1g and 1h). Pairwise comparisons found SED-C mice had lower energy expenditure compared to EXT-C (p < 0.0001, −3.9 kcal/kg body mass/h), SED-S (p < 0.0001, −3.8 kcal/kg body mass/h), and EXT-S mice (p < 0.0001, −5.0 kcal/kg body mass/h). EXT-S mice also had higher energy expenditure during the light cycle compared to SED-S mice, but this difference was not statistically significant (p = 0.08, 1.3 kcal/kg body mass/h) (Figure 1f & 1g). Similarly, pairwise comparisons found SED-C mice had lower energy expenditure compared to EXT-C (p < 0.0001, −3.4 kcal/kg body mass/h), SED-S (p < 0.0001, −4.4 kcal/kg body mass/h), and EXT-S mice (p < 0.0001, −4.3 kcal/kg body mass/h) (Figure 1f & 1h). There was no difference in EXT-C or EXT-S mice, suggesting there wasn’t an additive effect of sclerostin and exercise training on energy expenditure. This difference in energy expenditure may contribute to discrepancies in weight gain and food intake with sclerostin injection.

**Figure 1.**
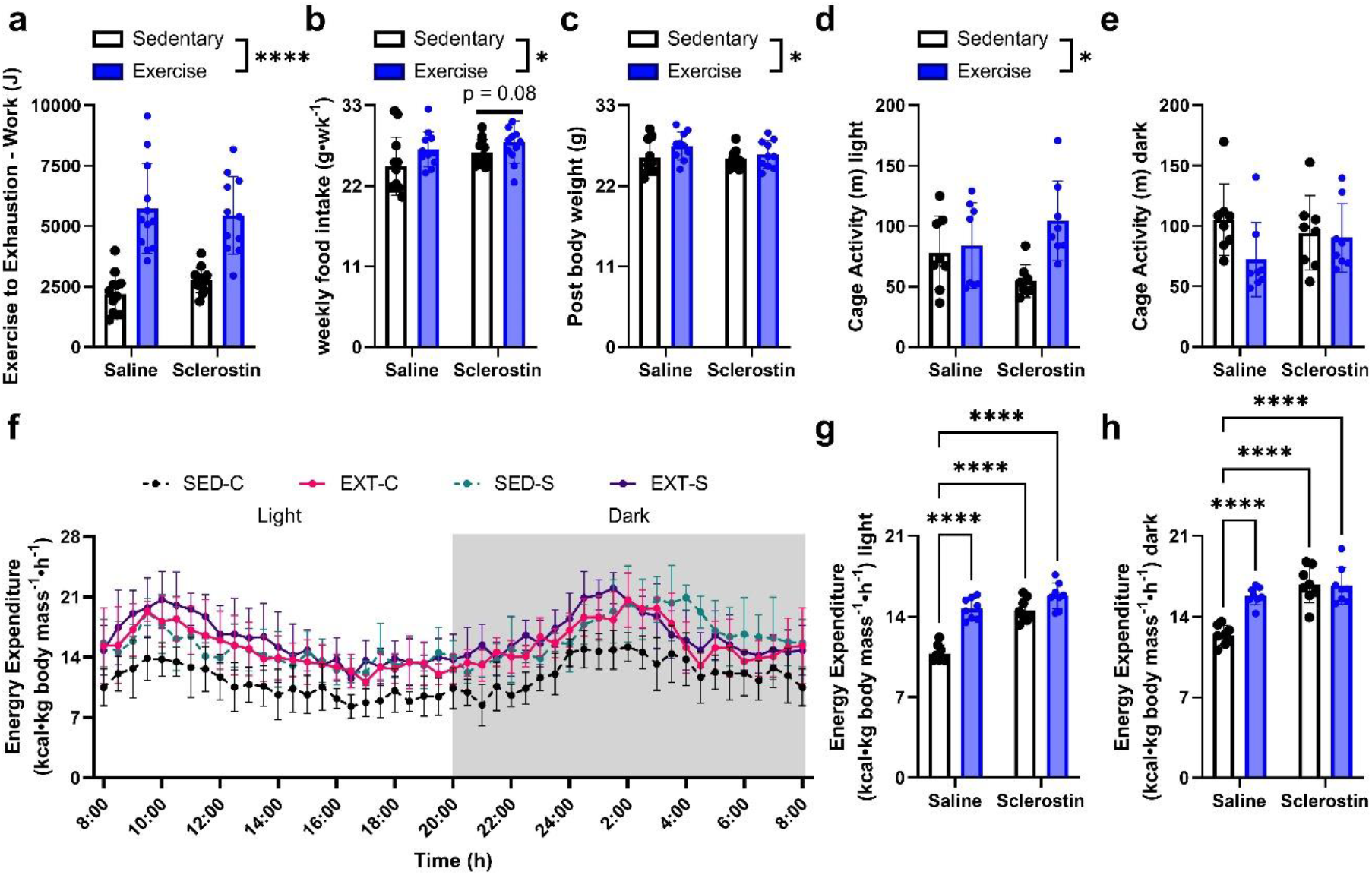
Sclerostin influences food intake and energy expenditure but does not further alter their responses to exercise training. (a) Work; (b) weekly food intake; (c) post-intervention body weight; cage activity during (d) light and (e) dark cycles; (f) 24h energy expenditure plotted at 30 min intervals, and 12h mean energy expenditure during (g) light and (h) dark cycles. Light and dark cycles are differentiated on figure ‘f’ by white and grey area, respectively. *(p<0.05) and ****(p<0.0001) indicate differences between specific groups or conditions (e.g., exercise vs. sedentary) (with black bars). A line over both columns of a specific injection group (e.g., sclerostin) indicate a group main effect. Data is presented as mean±SD. 2-way ANOVAs were used to assess exercise and injection main effects and following significant interactions pairwise comparisons were assessed using *Tukey* correction.

Exercise training is known to improve insulin sensitivity and influence resting fat and carbohydrate oxidation. Additionally, insulin sensitivity is improved in models of inhibited sclerostin action [18]. Assessment of resting blood glucose found it was higher in sclerostin injected mice compared to saline injected mice (p = 0.009, +0.26 mM) (Figure 2a). However, when AUC was assessed for the percent change in blood glucose across 1h following ITT, there was no difference between EXT and SED mice (main effect for exercise p = 0.5) or between sclerostin or saline injection (main effect for injection p = 0.1; exercise*injection interaction p = 0.6) (Figure 2b). Like resting levels, blood glucose concentration was again elevated at 60 min following ITT in sclerostin injected mice compared to saline (p = 0.03, +0.44 mM) (Figure 2a).

**Figure 2.**
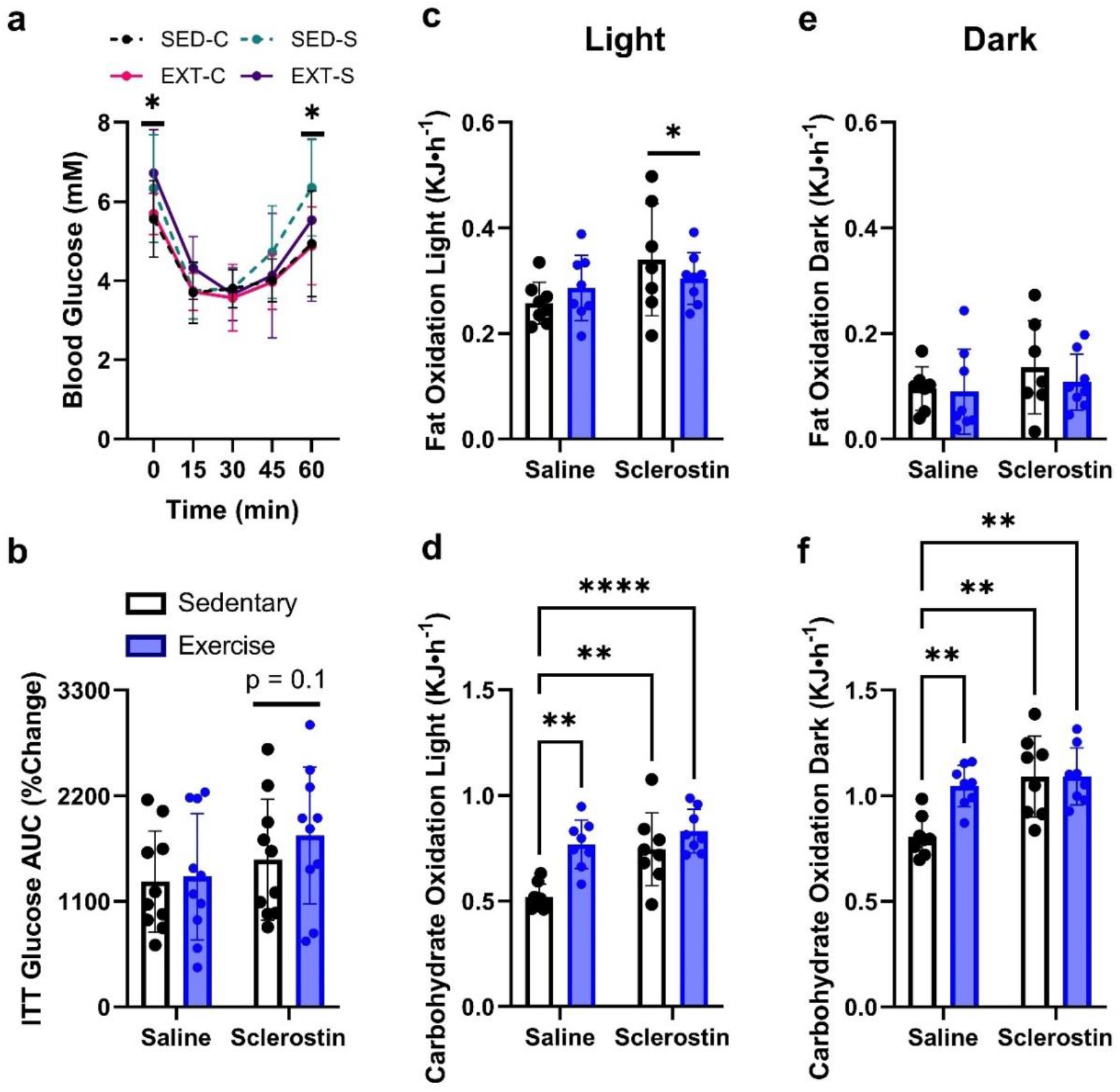
Sclerostin minimally influences insulin sensitivity and increases carbohydrate oxidation similarly to exercise training. (a) Blood glucose concentration across 60 min following ITT, (b) area under the curve (AUC) of the percent change of blood glucose to ITT, 12h mean fat oxidation and carbohydrate oxidation during (c and d) light and (e and f) dark cycles. *(p<0.05), **(p<0.01) and ****(p<0.0001) indicate differences between specific groups or conditions (e.g., exercise vs. sedentary) (with black bars). A line over columns or data points of a specific injection group (e.g., sclerostin) indicate a main effect for injection. Data is presented as mean±SD. 2-way ANOVAs were used to assess exercise and injection main effects and interactions and following significant interactions pairwise comparisons were assessed using *Tukey* correction.

To help explain differences in food intake and body mass in sclerostin injected mice compared to saline, we used VO_2_ and VCO_2_ data to determine fat and carbohydrate oxidation [26, 29]. Fat oxidation during the light cycle was not different between EXT and SED mice (main effect for exercise p = 0.9) but was higher in the sclerostin compared to saline injected mice (main effect for injection p = 0.048, +0.05 kJ/h) (Figure 2c). There was no difference in fat oxidation between EXT and SED mice (main effect for exercise p = 0.5) or sclerostin and saline injected mice (main effect for injection p = 0.3) during their dark cycle (Figure 2e). Carbohydrate oxidation during the light cycle was higher in EXT mice compared to SED mice (main effect for exercise p = 0.0004, grand mean difference = +0.17 kJ/h)) and higher in sclerostin injected mice compared to saline injected mice (main effect for injection p = 0.002, grand mean difference = +0.15 kJ/h). There was also a trend for an exercise*injection interaction (p = 0.06). Pairwise comparisons found SED-C mice had lower carbohydrate oxidation compared to EXT-C (p = 0.002, −0.25 kJ/h), SED-S (p = 0.004, −0.23 kJ/h), and EXT-S mice (p<0.0001, −0.31 kJ/h) (Figure 2d). Similarly, there was an exercise*injection interaction (p = 0.02) and simple effects for injection (p = 0.002, +0.17 kJ/h) and exercise (p = 0.02. +0.12 kJ/h) for carbohydrate oxidation during the dark cycle. Pairwise comparisons found SED-C mice had lower carbohydrate oxidation compared to EXT-C (p = 0.009, −0.24 kJ/h), SED-S (p = 0.002, −0.28 kJ/h), and EXT-S mice (p = 0.002, −0.28 kJ/h) (Figure 2f). These differences in fuel utilization/energy expenditure may help explain why there were contrasting findings in food intake (higher in sclerostin injected mice) and body weight (no difference).

### 3.2 Sclerostin does not prevent exercise-induced adaptations to bone

Given sclerostin is a potent inhibitor of bone formation [30], we next examined whole body bone mineral properties and tibial microarchitecture, a bone region known to demonstrate exercise induced responses as a result of increased mechanical loading (e.g., [31]). Whole body *in vivo* change in BMC was not influenced by exercise training (p = 0.1) or sclerostin injection (p = 0.6) and there was no exercise*injection interaction (p = 0.6) (Figure 3a). Whole body *in vivo* change in BMD was also not influenced by sclerostin injection (p = 0.3) or exercise training (p = 0.1) and there was no exercise*injection interaction (p = 0.2) (Figure 3b). Exercise training demonstrated improved tibia bone structure outcomes, characterized by greater BV/TV (main effect for exercise p < 0.0001, grand mean difference = +3.7) (Figure 3c and d), Tb.Th, and Tb.N, as well as greater Ct.Ar/Tt.Ar (main effect for exercise p = 0.01, grand mean difference = +1.3) and Ct.Th and decreased Ma.Ar in EXT mice compared to SED mice. However, sclerostin alone (p > 0.05) or in combination with exercise (p > 0.05) had no effect on trabecular or cortical bone outcomes at the proximal tibia and tibia midpoint, respectively (Table 1); with exception to main effects for sclerostin injection suggesting greater Tb.Sp (p = 0.03, grand mean difference = +0.012) and decreased Ct.Th when compared to saline injected mice.

**Figure 3.**
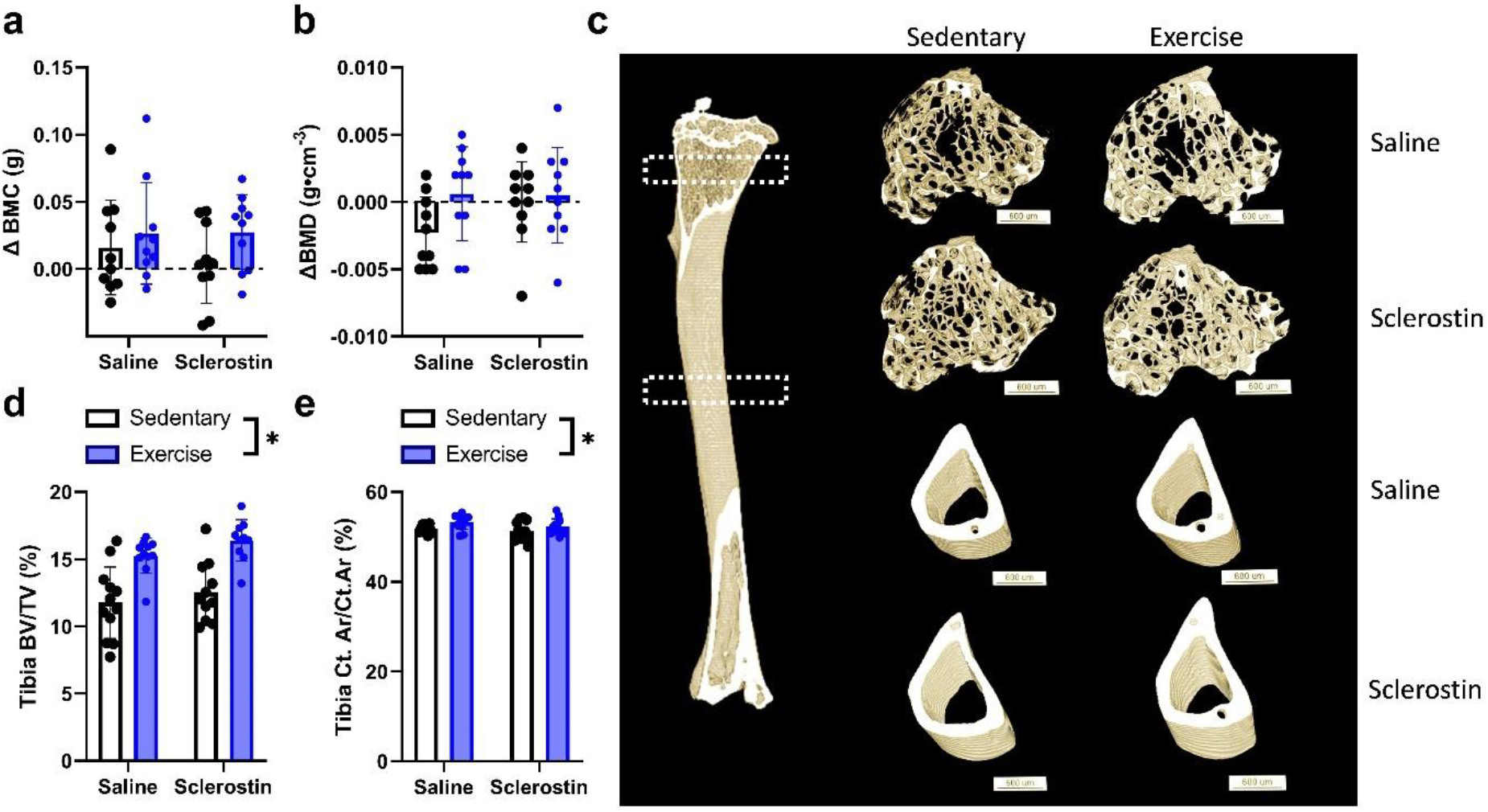
Sclerostin does not influence exercise training induced improvements in tibial microarchitecture. (a) Change in bone mineral content (BMC) from pre- to post-intervention. (b) change in bone mineral density (BMD) from pre- to post-intervention, (c) 3D representative images for endpoint trabecular (top 2 panels) and cortical bone (bottom 2 panels), (d) tibia trabecular BV/TV percentage, and € Ct Ar/Tt Ar percentage. *(p<0.05) indicate differences between specific groups or conditions (e.g., exercise vs. sedentary) (with black bars). Data is presented as mean±SD. 2-way ANOVAs were used to assess exercise and injection main effects and interactions and following significant interactions pairwise comparisons were assessed using *Tukey* correction.

**Table 1.** Trabecular and cortical bone parameters are enhanced with exercise training and are minimally impacted by 4 weeks of sclerostin injections.

|  | Saline |  | Sclerostin |  | 2-Way ANOVA |  |  |
| --- | --- | --- | --- | --- | --- | --- | --- |
|  | Sedentary | Exercise | Sedentary | Exercise | Exercise | Injection | Interaction |
| **Trabecular Bone Outcomes** |  |  |  |  |  |  |  |
| **BV/TV (%)** | 11.76±2.66 | 15.27±1.31 | 12.54±2.23 | 16.39±1.54 | <0.001 | 0.1 | 0.8 |
| **Tb.Th (mm)** | 0.044±0.025 | 0.052±0.003 | 0.046±0.004 | 0.052±0.003 | <0.001 | 0.3 | 0.6 |
| **Tb.Sp (mm)** | 0.21±0.025 | 0.20±0.015 | 0.20±0.023 | 0.19±0.017 | 0.08 | 0.03 | 0.3 |
| **Tb.N (1·mm<sup>-1</sup>)** | 2.65±0.43 | 2.98±0.24 | 2.79±0.36 | 3.12±0.33 | 0.002 | 0.2 | 1 |
| **DA (AU)** | 2.01±0.19 | 2.02±0.24 | 2.15±0.36 | 1.99±0.14 | 0.2 | 0.3 | 0.1 |
| **Conn.Dn (1·mm<sup>-3</sup>)** | 3.91E<sup>-4</sup>±5.71E<sup>-5</sup> | 3.66E<sup>-4</sup>±4.81E<sup>-5</sup> | 3.80E<sup>-4</sup>±5.49E<sup>-5</sup> | 4.02E<sup>-4</sup>±4.04E<sup>-5</sup> | 0.9 | 0.4 | 0.1 |
| **Cortical Bone Outcomes** |  |  |  |  |  |  |  |
| **Ct.Ar/Tt.Ar (%)** | 51.83±0.92 | 53.34±1.65 | 51.27±2.13 | 52.28±1.77 | 0.01 | 0.1 | 0.6 |
| **Ps.Pm (mm)** | 7.70±0.46 | 7.80±0.33 | 7.66±0.46 | 7.95±0.43 | 0.1 | 0.6 | 0.4 |
| **Ec.Pm (mm)** | 3.12±0.22 | 3.21±0.24 | 3.10±0.26 | 3.25±0.23 | 0.1 | 0.9 | 0.7 |
| **Ma.Ar (mm<sup>2</sup>)** | 0.63±0.058 | 0.61±0.069 | 0.62±0.058 | 0.63±0.065 | 0.04 | 0.3 | 0.9 |
| **Ecc (AU)** | 0.67±0.026 | 0.69±0.040 | 0.68±0.038 | 0.70±0.027 | 1 | 0.9 | 0.5 |
| **Ct.Th (mm)** | 0.172±0.006 | 0.176±0.004 | 0.169±0.006 | 0.173±0.004 | 0.01 | 0.03 | 0.9 |

BV/TV = bone volume fraction; Tb.Th = trabecular thickness; Tb.Sp = trabecular separation; Tb.N = Trabecular number; DA = degree of anisotropy; Conn.Dn = connectivity density. Ct.Ar/Tt.Ar = cortical area fraction; Ps.Pm = periosteal perimeter; Ec.Pm = endocortical perimeter; Ma.Ar = medullary area; Ecc = eccentricity; Ct.Th = cortical thickness. Data is presented as mean±SD. 2-way ANOVAs were used to assess exercise and injection main effects and interactions.

### 3.3 Sclerostin prevents exercise training induced improvements in lean mass and subcutaneous fat mass

Given peripheral sclerostin inhibition with neutralizing antibodies can reduce fat mass [18], we also examined if sclerostin injections could prevent exercise induced adaptations in body composition, fat pad mass, and WAT adipocyte cell size. We found an exercise*injection interaction for change in lean mass (p = 0.03). This was the result of EXT-C mice having a larger increase in lean mass compared to SED-C (p = 0.002, +2.2g) and SED-S mice (p = 0.005, +1.97g) and trended to be higher than EXT-S mice (p = 0.06, +1.45g) (Figure 4a). There was also an exercise*injection interaction for change in body fat percentage (p = 0.02). This was the result of EXT-C decreasing their body fat percentage across the study and SED-C increasing their fat mass (p = 0.006, +1.96%) while SED-S and EXT-S mice had no change in fat mass (p > 0.9) (Figure 4b). eWAT necropsy weight was also lower in EXT mice compared to SED mice (p = 0.0004, −3.1 mg). However, sclerostin did not influence this adaptation in EXT mice (exercise*injection interaction: p = 0.4) or in general (main effect for injection: p = 0.6) (Figure 4c). There was an exercise*injection interaction for iWAT necropsy weight (p = 0.049) due to EXT-C mice having a lower iWAT mass compared to SED-C (p = 0.02, −1.86 mg) with no difference in SED-S and EXT-S iWAT mass (p > 0.9) (Figure 4d). iBAT mass was not different between EXT and SED mice (p = 0.2) or sclerostin and saline injected mice (p = 0.4) and there was no exercise*injection interaction (p = 0.1) (Figure 4e). eWAT adipocyte cross sectional area was not different between EXT and SED mice (p = 0.3) or sclerostin and saline injected mice (p = 0.09) and there was no exercise*injection interaction (p = 0.3) (Figure 4f and 4i). In contrast, iWAT adipocyte cross sectional area had an exercise*injection interaction (p = 0.005). This was a result of EXT-C mice having a lower iWAT mean adipocyte cross sectional area compared to SED-C mice (p = 0.02, −234 µm^2^) while there was no difference between SED-S and EXT-S (p = 0.6) (Figure 4g and 4i). Similarly, there was an exercise*injection interaction for percentage of multilocular adipocytes in iWAT (p = 0.03). While there was no significant pairwise comparisons or trends (i.e., p<0.1), this interaction was a result of EXT-C mice tending to have a higher percentage of multilocular adipocytes in iWAT compared to SED-C mice (p = 0.17, +3.2%) while SED-S mice tended to have a higher percentage compared to EXT-S mice (p = 0.7, +1.7%) (Figure 4h and 4i). Together, it appears as though increasing sclerostin prevents exercise training induced adaptations in body composition, particularly in subcutaneous iWAT.

**Figure 4.**
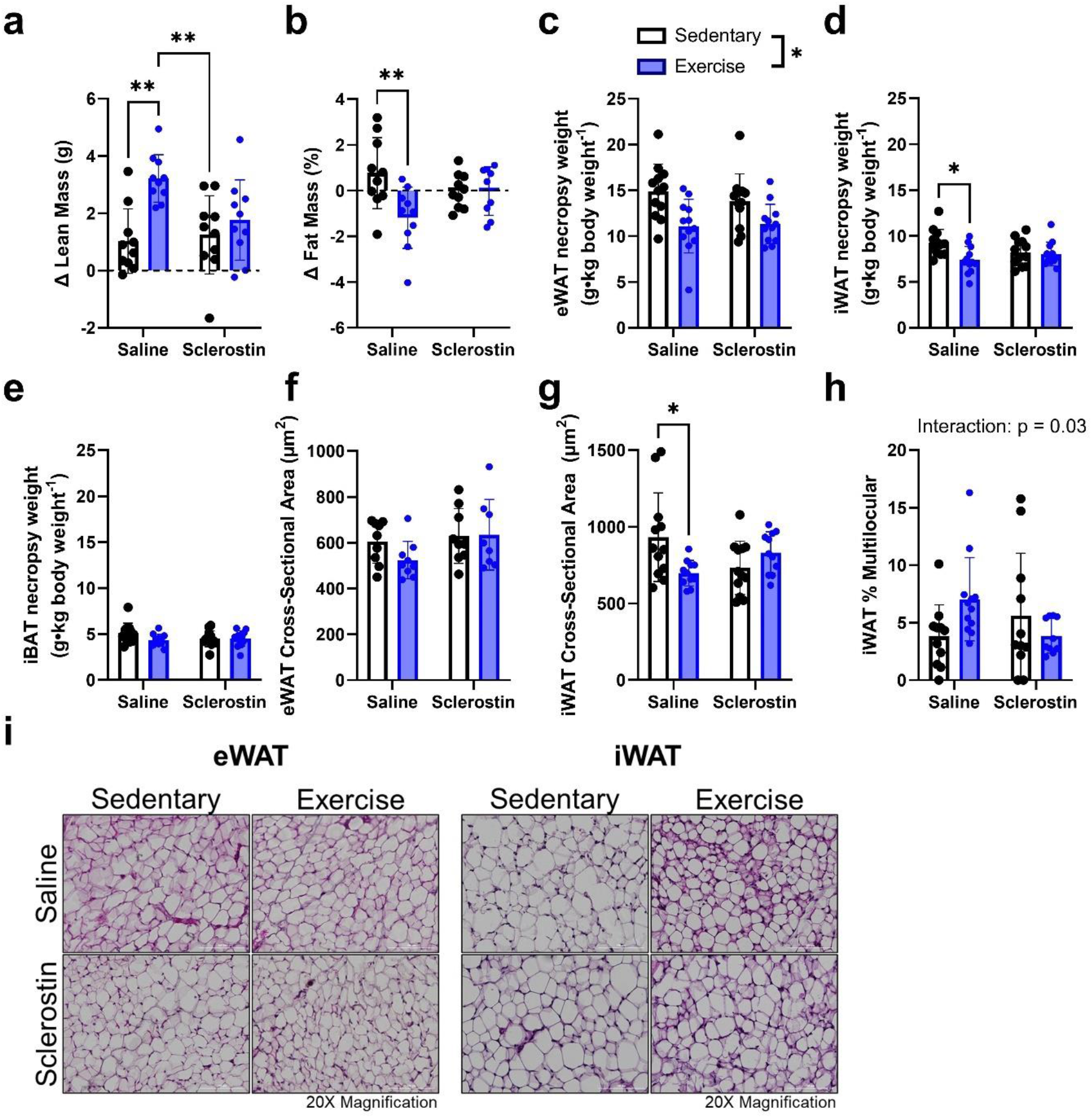
Sclerostin prevents exercise training induced improvements in body composition and iWAT morphology. (a) Change in lean mass from pre- to post-intervention, (b) change in fat mass from pre- to post-intervention, necropsy weight for (c) eWAT, (d) iWAT, and (e) iBAT, (f) eWAT adipocyte cross-sectional area, (g) iWAT adipocyte cross-sectional area, (h) iWAT percentage of multilocular cells, and (i) representative histological samples for eWAT and iWAT. *(p<0.05) and **(p<0.01) indicate differences between specific groups or conditions (e.g., exercise vs. sedentary) (with black bars). Significant exercise*injection interactions with no significant pairwise comparisons are presented in text on graphs. Data is presented as mean±SD. 2-way factorial ANOVAs were used and following significant interactions pairwise comparisons were assessed using Tukey correction.

Given sclerostin influenced exercise training induced adaptations to iWAT adipocyte cross sectional area and the number of multilocular cells we also assessed markers of lipolysis (Figure 5). There was no effect of either exercise training or sclerostin injection on the activation phosphorylation status of either hormone sensitive lipase (HSL) (Figure 5a and b) or adipose triglyceride lipase (ATGL) (Figure 5c).

**Figure 5.**
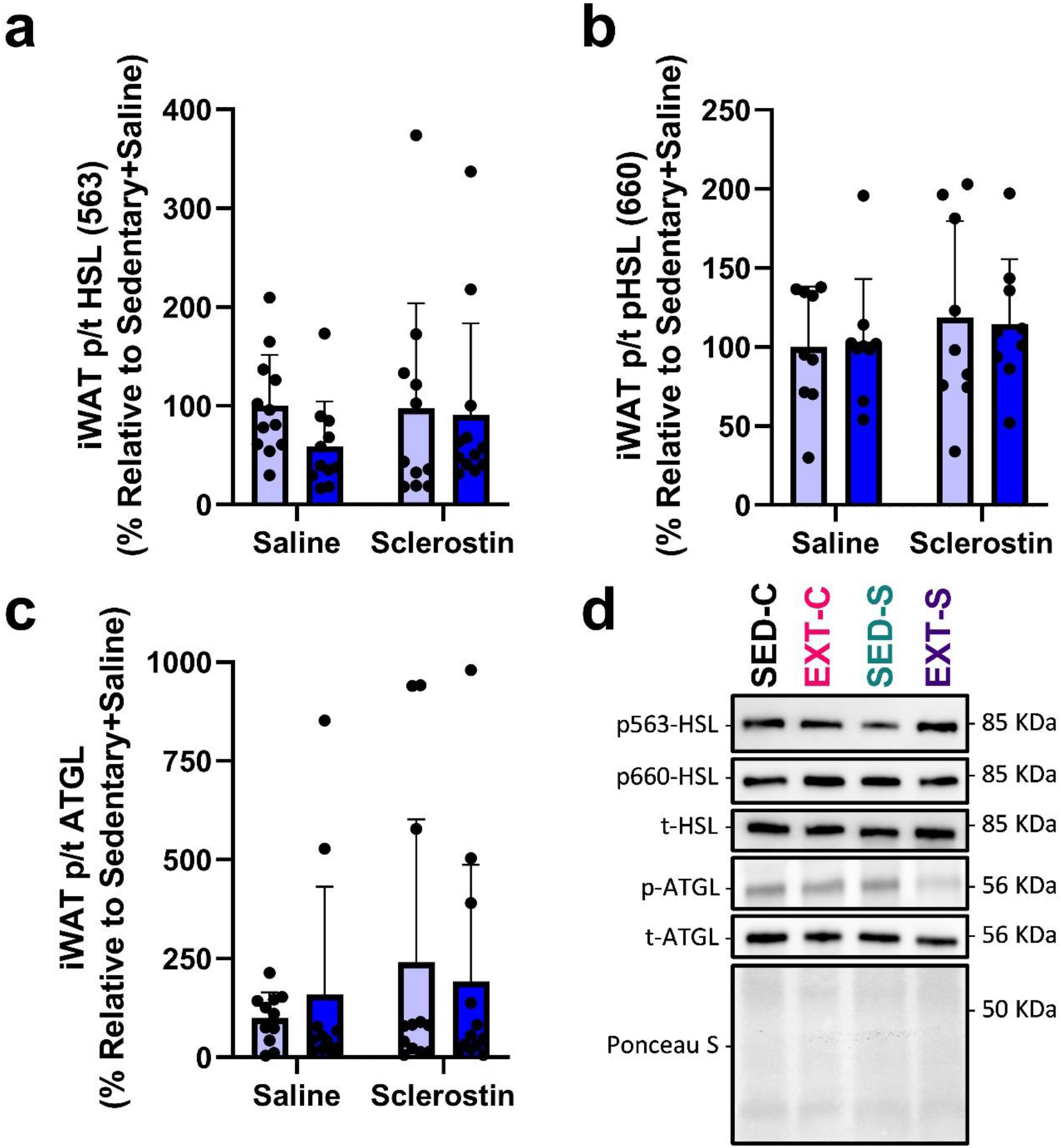
Exercise training and sclerostin injections do not influence the resting activation status of lipolysis enzymes in iWAT. iWAT phosphorylation status of (a and b) HSL and (c) ATGL and (d) their representative immunoblots. Data is presented as mean±SD. 2-way factorial ANOVAs were used. HSL = hormone sensitive lipase; ATGL = adipose triglyceride lipase.

Sclerostin inhibits Wnt/β-catenin signalling, a pathway that can regulate mitochondrial biogenesis and oxidative capacity [32]. Additionally, exercise training has been shown to augment subcutaneous WAT Wnt signalling in mice [21, 33] and humans [11]. Therefore, we next examined serum sclerostin concentration and iWAT sclerostin content, Wnt/β-catenin signalling, and markers of mitochondrial content and biogenesis. As expected, serum sclerostin was higher in sclerostin injected mice compared to saline (p = 0.0004, +101.0 pg/ml) and was not different in EXT compared to SED mice (p = 0.1, −42.2 pg/ml) and there was no exercise*injection interaction (p = 0.1) (Figure 6a). The magnitude of change in peripheral sclerostin with daily treatment of recombinant sclerostin is consistent to increases observed in humans with metabolic syndrome/insulin resistance compared to controls [34, 35], but slightly less of an increase compared to mice that have HFD (10-week intervention) induced obesity and perturbed glucose homeostasis/insulin resistance [18]. iWAT sclerostin was also higher in sclerostin injected mice compared to saline (p < 0.0001, +77.9%). Like what we have seen in humans [11], iWAT sclerostin trended to be lower in EXT compared to SED mice (p = 0.068, - 23.7%) and there was no exercise*injection interaction (p = 0.5) (Figure 5b and 5g). Despite increased iWAT sclerostin, iWAT ser9 p/t GSK3β content was higher in sclerostin injected mice compared to saline (p < 0.0001, +738.9%) and was higher in EXT mice compared to SED mice (p = 0.004, +299.1%) and there was no exercise*injection interaction for iWAT sclerostin (p = 0.3) (Figure 6e and 6g). iWAT Lrp4 content had an exercise*injection interaction which was a result of Lrp4 being higher in EXT-C mice compared to SED-C (p = 0.005, +290%) while EXT-S mice were not different compared to SED-C (p > 0.9, −43%) or SED-S mice (p > 0.9, −23%) (Figure 6c and 6g). iWAT β-catenin content was also higher in EXT mice compared to SED mice (main effect for exercise p = 0.03, grand mean difference = +231.9%), which matches responses we have seen in humans [11], however, this response was not different in sclerostin injected mice compared to saline (p = 0.2) and there was no exercise*injection interaction (p = 0.9) (Figure 6f and 6g) despite sclerostin being a known Wnt/β-catenin inhibiter. Wnt signalling is known to regulate mitochondrial biogenesis, and again there was no effect of sclerostin injection on any markers of mitochondrial content that increased with exercise training within iWAT (e.g., CI-NDUFB8, CIV-MTO1, and CV-ATP5A) (Figure 6h and 6g).

**Figure 6.**
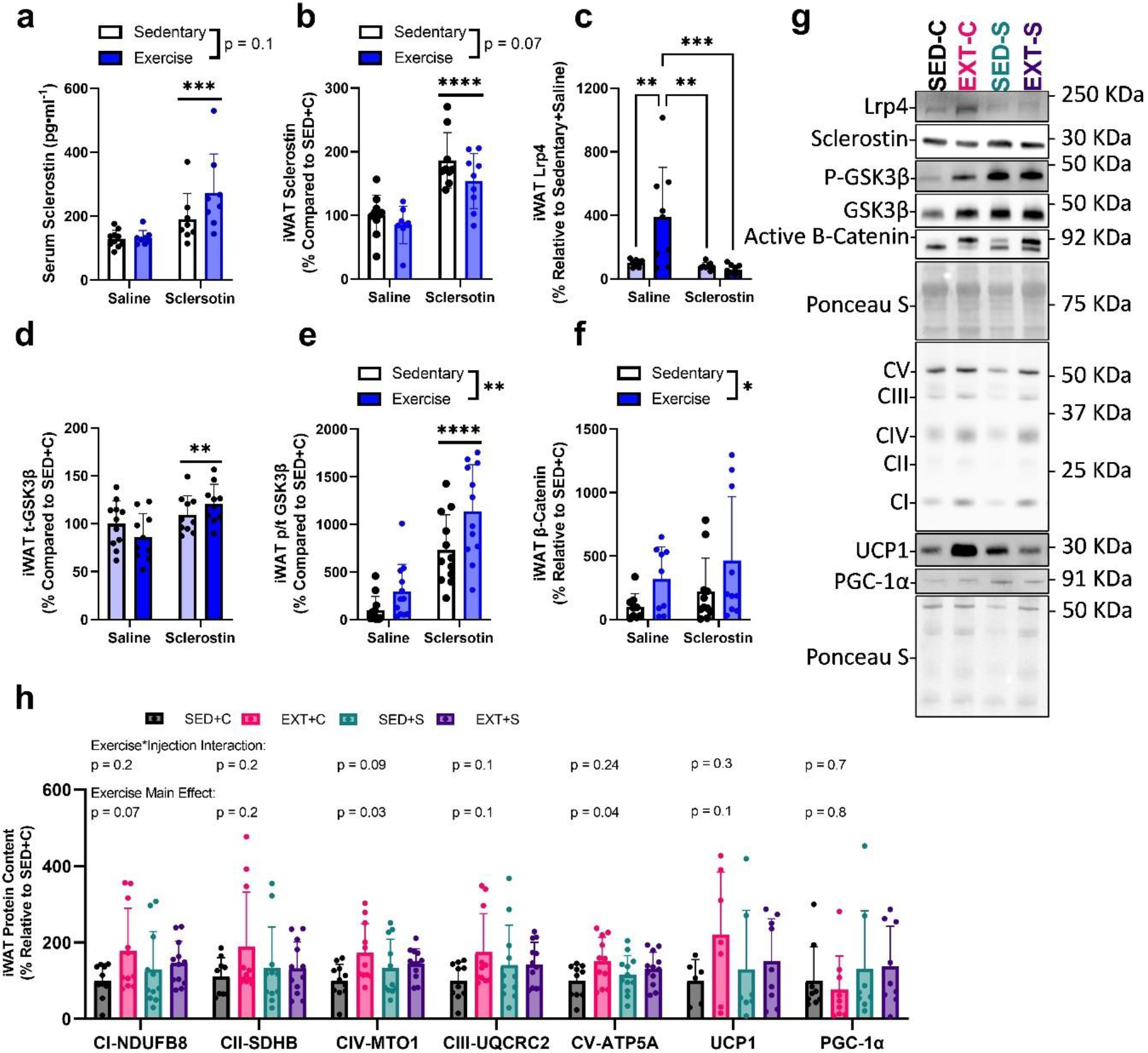
Sclerostin injections inhibited exercise training induced increase in iWAT Lrp4 but did not influence Wnt/β-catenin signalling or markers of mitochondrial content. (a) Serum sclerostin and iWAT (b) sclerostin, (c) Lrp4, (d) t-GSK3β, (e) p/t GSK3β, (f) β-catenin, (h) markers of oxidative phosphorylation/ mitochondrial biogenesis, and (g) their representative immunoblots. *(p<0.05), **(p<0.01), ***(p<0.001), and ****(p<0.0001) indicate differences between specific groups or conditions (e.g., exercise vs. sedentary) (with black bars). A line over both columns of a specific injection group (e.g., sclerostin) indicate a group main effect. For figure ‘h’ exercise*injection interaction and exercise main effect p-values for specific proteins are directly above them. There were no significant injection effects, and these values are reported in text. Data is presented as mean±SD. 2-way factorial ANOVAs were used and following significant interactions pairwise comparisons were assessed using *Tukey* correction. CI-NDUFB8 = NADH dehydrogenase [ubiquinone] 1 Beta Subcomplex Subunit; CII-SDHB = Succinate Dehydrogenase (ubiquinone) Iron-Sulfur Subunit; CIV-MTO1 = Mitochondrial Translation Optimization Factor 1; CIII-UQCR2 = Cytochrome b-c1 Complex Subunit 2; CV-ATP5A = ATP Synthase F1 Subunit Alpha; UCP1 = uncoupling protein 1; PGC-1α = The Peroxisome Proliferator-Activated Receptor Gamma Coactivator 1-Alpha.

## 4 Discussion

Exercise training has beneficial effects on metabolism (glucose handling and fat oxidation) and tissue remodelling (e.g., musculoskeletal system). Recent studies have shown that proteins secreted during acute exercise and those that are regulated at rest with exercise training can influence these adaptations [36]. In this study, we provide evidence for a potential role of sclerostin in regulating exercise training induced adaptations to body composition. Specifically, we re-confirm that sclerostin is reduced (trend) and Wnt/β-catenin signalling is enhanced in subcutaneous WAT following exercise training [11]. However, we did not observe evidence for exercise training induced reduction in peripheral sclerostin that has been shown in humans. When mice were injected with recombinant sclerostin to prevent this reduction, EXT induced increases in body mass, lean mass, and iWAT percent multilocular cells and reductions in iWAT fat pad mass and adipocyte cell size were prevented. There was an exercise training induced reduction in eWAT fat pad mass and no influence of sclerostin on this depot’s mass or adipocyte cell size, suggesting depot specific effects of sclerostin. Exercise training also increased energy expenditure and improved tibial bone structure while sclerostin did not influence these responses. However, sclerostin injection alone increased energy expenditure and had minimal effects on tibial microarchitecture. Our findings suggest small changes in sclerostin’s concentration (i.e., following EXT) can influence body composition, tissue morphology (iWAT), and metabolism.

This study found no impact of sclerostin injection on exercise training induced adaptations in whole body bone mineral properties or tibia microarchitecture. While this is likely due to the short duration of the study or the exercise stimulus being more potent than the catabolic effect of the sclerostin dose, it may suggest circulating sclerostin is related to its effects on tissues beyond bone (e.g., WAT or lean mass). This would indicate more profound endocrine action of sclerostin compared to its autocrine/paracrine action within bone. This could explain why we see a similar bone turnover response despite a larger and more sustained increase in circulating sclerostin to acute exercise in humans that have high adiposity compared to lean controls [12]. We speculate the influence of exercise training and adiposity on WAT sclerostin content and the response to acute exercise is due to alterations in protein content within the bone multicellular unit. This is likely due to the multiple bouts of acute exercise acting as a model of mechanical shear stress, which has been shown to increase lysosomal degradation of sclerostin *in vitro* [37]. Indeed, multiple bouts of acute exercise (i.e., exercise training) reduces bone sclerostin expression in mice [13–16] and may contribute to reductions in resting serum sclerostin concentration [6–10] and scWAT sclerostin content [11] over time.

Previous studies have shown sclerostin impacts visceral and subcutaneous fat mass through the regulation of adipocyte metabolism, ultimately, influencing cell size and differentiation *in vivo* [1, 5, 18]. Specifically, when sclerostin action is inhibited, there is an increase in insulin sensitivity and fat oxidation, reduction in both visceral and scWAT adipocyte cell size, as well as protection against obesogenic diet induced liver steatosis, insulin resistance, and fat mass expansion [18]. Adenovirus overexpression of sclerostin (+65% increase in circulating sclerostin) in mice also leads to a ∼2% reduction in lean mass, ∼3% increase in fat mass (equal increase in gonadal, inguinal, and retroperitoneal fat pad weight), and increased adipocyte hypertrophy [18]. These effects were accompanied by reduced expression of Wnt target genes and an increase in enzymes involved in de novo lipid synthesis and storage within WAT. Our findings are in line with sclerostin reducing insulin sensitivity (higher basal and 60 min post-ITT blood glucose) and reductions in scWAT and lean mass, however, our results do not support an effect of sclerostin on visceral WAT and contrast the effect on fat oxidation. This may be due to differences in mechanism of action (recombinant protein compared to adenovirus overexpression), dose, length of treatment, or route of administration (e.g., continuous elevation through hepatic expression vs. frequent injections) between studies. EXT mice had reductions in iWAT sclerostin, and our low dose appears to sufficiently inhibit associated adaptations in this depot. Visceral WAT may be less responsive to sclerostin treatment, and a larger dose may be required (e.g., low receptor content). Our model also found sclerostin increased energy expenditure and carbohydrate and fat oxidation while others have found no effect on energy expenditure [18]. This alteration in metabolism appears to be related to changes observed in food intake (sclerostin increased food intake) and body weight (no effect of sclerostin). The difference in body composition between EXT control and sclerostin injected mice also suggests a potential catabolic state in the sclerostin injected mice. Future studies are needed to assess which mechanisms are influencing this change in substrate utilization and the influence on body composition. We did find EXT increased iWAT Lrp4 content, sclerostin’s receptor, which was lost with sclerostin injection. This may be a desensitization response due to an overstimulation with sclerostin in the EXT-S group. Lrp4 has been extensively studied in muscle and brain and has a critical role in neuromuscular junction formation and acetylcholine receptor activation [38–42]. While there is only one study examining Lrp4’s role in regulating adipose tissue metabolism and mass [5], exercise appears to regulate its content and sclerostin can inhibit this adaptation. This may explain some of the adaptations in adipose tissue mass and morphology, given Lrp4 knockout mice have reduced WAT mass.

Given the influence of sclerostin treatment on EXT induced adaptations in iWAT mass and histology, we further examined this depot for changes in cell signalling, markers of mitochondrial content, and lipolysis markers. EXT mice had higher markers of mitochondrial content that was not affected by sclerostin injections. This contradicted our histological data (reduced adipocyte cell size and increased multilocular cells) and previous studies that have found inhibited Wnt signalling promotes adipocyte expansion and inhibits adipocyte mitochondrial biogenesis [32]. However, Wnt/β-catenin signalling has distinct roles in regulating mature adipocyte metabolism and function depending on the model and component examined. Specifically, adipocyte β-catenin knockout models do not result in altered fat mass or metabolism in chow fed mice, but are protected against diet induced obesity by preventing lipogenic gene expression, de novo lipogenesis, and lipid desaturation [43], particularly in subcutaneous WAT [44]. In contrast, Wnt overexpression within adipocytes is protective against obesity by promoting oxidative metabolism and improving insulin sensitivity/glucose tolerance [32, 45, 46]. These lines of evidence point to a discrete effect of individual components of Wnt signal transduction in regulating fat mass expansion in response to HFD feeding. While we observed apparent inhibition of GSK3β via serine 9 phosphorylation with EXT, surprisingly, sclerostin injections also increased its phosphorylation. While GSK3β has been shown to inhibit the thermogenic gene program [47], we found no response in UCP1 or PGC-1α content to EXT or sclerostin injections. Additionally, active β-catenin content (i.e., unphosphorylated) was increased with EXT and there was no effect of sclerostin injection. Together these findings suggest there may be compensation due to reduction in sclerostin receptor content (Lrp4), connecting pathways [48], or neighbouring cells [43, 49] to try and prevent the Wnt inhibitory effect of our continuous sclerostin injections. It is also important to note that these signalling events are temporal and may not reflect the dynamic changes in cell signalling that occur shortly after injection which are likely driving morphological changes. Future studies should assess the acute effect of sclerostin injections *in vivo* on Wnt/β-catenin signalling across tissues at several time points post injection.

This study suggests iWAT is the most influenced fat depot by exercise induced sclerostin. iWAT controls peripheral tissue homeostasis through the storage and timely release of substrates that contributes to energy expenditure in peripheral tissues (e.g., liver, and skeletal muscle) [21, 50, 51]. However, the discrepancy in energy expenditure and carbohydrate and fat oxidation between sclerostin and saline injected mice suggests other metabolically active tissues, like bone [52, 53], brain [54], liver [18, 55], and skeletal muscle [56] may be either directly (i.e., sclerostin binding and influencing cell differentiation and metabolism) or indirectly (i.e., influence on lipolysis in iWAT) contributing to these changes in metabolism. While we found no effect of EXT or sclerostin injection on activation of lipolytic enzymes, it may be because we measured basal levels and not their activation in response to physiological stimuli (e.g., adrenergic activation) that could reveal functional differences. Wnt signalling within skeletal muscle has been shown to cause a shift to a oxidative fibre type [57, 58] as well as the regulation of thermogenesis, which may also explain sclerostin induced differences in energy expenditure, fat oxidation, and carbohydrate oxidation observed in this study [59]. Additionally, given our data on Lrp4 in adipose tissue future studies should assess the effect increasing peripheral sclerostin would have on Lrp4 content in skeletal muscle and the subsequent influence on fibre type and neuromuscular junction integrity known to be important for muscle strength [40].

## 5 Conclusion

These findings suggest a role of sclerostin/Lrp4 in regulating body composition, particularly scWAT, in response to exercise training. Our findings also support a role of sclerostin in regulating peripheral tissue metabolism, implicating it in the pathophysiology of conditions with elevated sclerostin (e.g., obesity, pre-diabetes, low energy availability).

## 6 Acknowledgements

This study was funded by a Natural Sciences and Engineering Research Council of Canada (NSERC) grant to P. Klentrou (grant # 2020-00014). N. Kurgan holds a doctoral Queen Elizabeth II Graduate Scholarship in Science and Technology (QEII-GSST).

